# *Shh* and ZRS enhancer co-localisation is specific to the zone of polarizing activity

**DOI:** 10.1101/050849

**Authors:** Iain Williamson, Laura A. Lettice, Robert E. Hill, Wendy A. Bickmore

**Affiliations:** MRC Human Genetics Unit, MRC Institute of Genetics and Molecular Medicine, Crewe Road, Edinburgh EH4 2XU, UK

**Keywords:** super-resolution, FISH, 5C, limb development, enhancer

## Abstract

Limb-specific *Shh* expression is regulated by the (~1 Mb distant) ZRS enhancer. In the mouse, limb bud restricted spatiotemporal expression of *Shh* occurs from ~E10-E11.5 at the distal posterior margin and is essential for correct autopod formation. Here, we have analysed the higher-order chromatin conformation of *Shh* in expressing and non-expressing tissues, both by fluorescence in situ hybridisation (FISH) and by chromosome conformation capture (5C). Conventional and super-resolution light microscopy identified significantly elevated frequencies of *Shh*/ZRS co-localisation only in the *Shh* expressing regions of the limb bud. However, *Shh*-ZRS spatial distances were consistently shorter than intervening distances to a neural enhancer in all tissues and developmental stages analysed. 5C identified a topologically associating domain (TAD) over the *Shh*/ZRS genomic region and enriched interactions between *Shh* and ZRS throughout E11.5 embryos. *Shh*/ZRS co-localisation, therefore, correlates with the spatiotemporal domain of limb bud-specific *Shh* expression, but close *Shh*/ZRS proximity in the nucleus occurs regardless of whether the gene or enhancer is active. We suggest that this constrained chromatin configuration optimises the opportunity for the active enhancer to locate and instigate *Shh* expression.

## Introduction

Chromatin-looping is a popular model by which very long-range enhancers can communicate with their target gene promoter (Benabdallah and Bickmore, 2015), however the relationship of loop formation and gene activation remains unclear. It has been suggested that enhancer-target gene contacts are preformed and present in tissues even where the target gene is not activated (Montavon et al., 2011; Ghavi-Helm et al., 2014). However, other reports indicate enhancer-gene looping is spatially and temporally restricted to cells where the target gene is active. This includes in the developing mouse limb, where elevated levels of co-localisation of the global control region (GCR) and its target 5′HoxD genes is only seen in the cells of the distal posterior portion of the E10.5 limb bud (Williamson et al., 2012).

The complex spatiotemporal circuit of gene regulation in the developing limb is a rich system in which to study the activity of distal regulatory elements and their mechanisms of action. The sonic hedgehog gene (*Shh*), encodes a morphogen that directs cell fate during organogenesis. Limb-specific expression of *Shh* is regulated by the ZRS enhancer positioned within an intron of *Lmbr1* ~1 Mb away at the opposite end of a large gene desert (Lettice et al., 2002; Lettice et al., 2003) (Figure 1A). The ZRS has a functional role in directing spatiotemporal *Shh* expression restricted to a region of the distal posterior mesenchyme of the limb bud known as the zone of polarizing activity (ZPA) (Saunders and Gasseling, 1968; Riddle et al., 1993). Limb-specific *Shh* expression is abrogated upon deletion of ZRS (Sagai et al., 2005), whereas point mutations across the 780-bp conserved sequence of the enhancer can induce anterior, ectopic *Shh* expression and can cause preaxial polydactyly (Lettice et al., 2003; Sagai et al., 2004; Lettice et al., 2008), triphalangial thumb (Furniss et al., 2008) or Werner mesomelic syndrome (VanderMeer et al., 2014).

**Figure 1.**
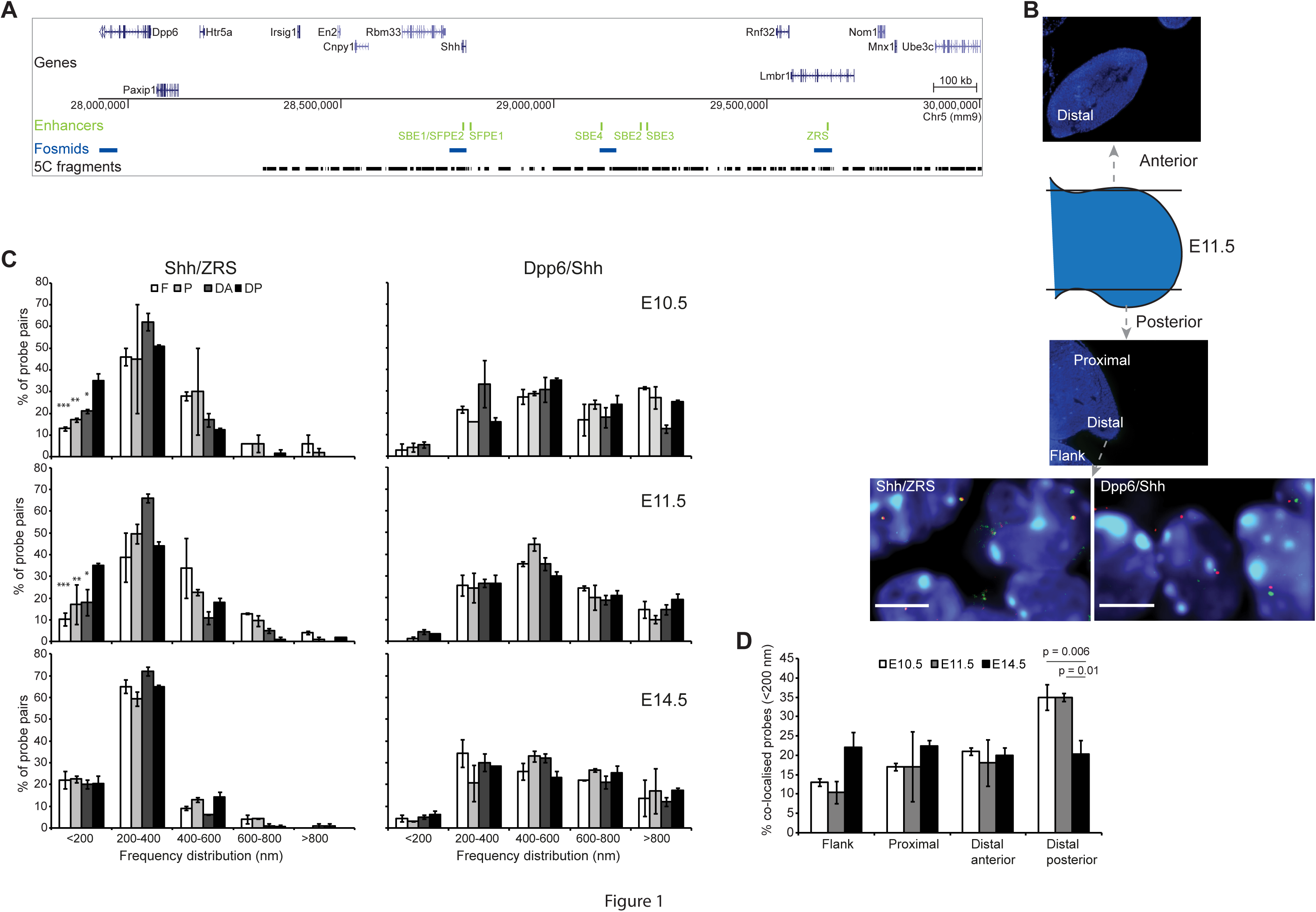
ZRS*Shh* proximity in the ZPA at E10.5 and E11.5. (**A**) (Top) Location of genes over a 2 Mb murine genomic locus containing *Shh*, with the position of tissue-specific *Shh* enhancers shown below in green. The bottom two tracks show the locations to which the fosmid probes used for FISH hybridize (blue) and the 3C fragments amplified for 5C (black). (**B**) Schematic indicating the position and plane of the tissue sections taken through the anterior and posterior parts of the E11.5 forelimb bud. Distal and proximal parts of the posterior limb bud and the distal anterior limb bud are shown, as is the flank mesoderm. Below are images of nuclei from E11.5 ZPA tissue sections showing *Shh*/ZRS and *Shh/Dpp6* probe pairs. Scale bars = 5μm. (**C**) Frequency distributions of FISH inter-probe distances (d) in 200 nm bins, between *Shh* and ZRS (left column), or *Shh* and *Dpp6* probes (right column) in proximal and distal regions of the murine forelimb bud and adjacent flank at E10.5, E11.5 and E14.5. F flank, P: proximal limb, DA: distal anterior limb, DP: distal posterior limb (ZPA in E10.5 and E11.5 sections). n = 70-130 (alleles). Error bars represent SEM obtained from two or three different tissue sections from 1-2 embryos. The statistical significance between data sets was examined by Fisher’s Exact Tests: * *p* < 0.05, ** *p* < 0.01, *** *p* < 0.001. (**D**) Comparison of the proportion of co-localised *Shh*/ZRS probe pairs (<200nm) across the three temporal developmental stages for distal anterior and posterior and proximal forelimb tissue and flank tissue. Error bars represent SEM obtained from two or three different tissue sections. The statistical significance between data sets was examined by Fisher’s Exact Tests.

Previously, fluorescence in situ hybridisation (FISH) and chromosome conformation capture (3C) (Amano et al., 2009) have reported increased associations between *Shh* and ZRS in E10.5 limb buds compared with other tissues. However, no significant difference in gene/enhancer co-localisation was detected between the ZPA and distal anterior tissue – where *Shh* is not normally expressed, or indeed in ZPA cells between wild-type and embryos with a deletion of the ZRS. This is consistent with a model of pre-formed enhancer-gene contacts. In contrast, FISH has revealed a significant decrease in *Shh*/ZRS co-localisation in E11.5 ZPA tissue from mouse embryos with a ZRS mutation which decreases ZRS long-range activity (Lettice et al., 2014), suggesting that ZRS/*Shh* juxtaposition is directly linked to *Shh* activation.

We have previously combined FISH and 3C carbon copy (5C) to elucidate the role of chromatin conformation in the long-range regulation of the 5’ *Hoxd* genes during distal limb bud development (Williamson et al., 2012; Williamson et al., 2014). Here we combined these methods to characterise the higher-order chromatin landscape over the *Shh* locus in tissue sections – including those derived from three discrete developmental stages of mouse limb bud development. Spatial proximity of *Shh* and ZRS, as inferred indirectly from enriched 5C interactions, was identified throughout E11.5 embryos, and 5C data confirmed that *Shh* and its known enhancers form a compact regulatory chromatin domain. However, using super-resolution microscopy we show that, despite *Shh* and ZRS being proximal to one another in the nucleus in all tissue types and temporal stages analysed, high levels of *Shh*/ZRS co-localisation occurs only in ZPA cells at the time of *Shh* activation.

## Materials and Methods

### FISH

For 3D FISH, E10.5, E11.5 and E14.5 embryos from CD1 mice were collected, fixed, embedded, sectioned and processed as previously described (Morey et al., 2007), except that sections were cut at 6 μm. Fosmid clones (Figure 1A, Table S1) were prepared and labelled as previously described (Morey et al. 2007). Between 160-240 ng of biotin-and digoxigenin-labeled fosmid probes were used per slide, with 16-24 μg of mouse Cot1 DNA (Invitrogen) and 10 μg salmon sperm DNA.

### Image analysis

For 3D analysis of tissue sections by conventional microscopy, slides were imaged with a Hamamatsu Orca AG CCD camera (Hamamatsu Photonics (UK) Ltd, Welwyn Garden City, UK), Zeiss Axioplan II fluorescence microscope with Plan-neofluar or Plan apochromat objectives, a Lumen 200W metal halide light source (Prior Scientific Instruments, Cambridge, UK) and Chroma #89014ET single excitation and emission filters (Chroma Technology Corp., Rockingham, VT) with the excitation and emission filters installed in Prior motorised filter wheels. A piezoelectrically driven objective mount (PIFOC model P-721, Physik Instrumente GmbH & Co, Karlsruhe) was used to control movement in the z dimension. Hardware control, image capture and analysis were performed using Volocity (Perkinelmer Inc, Waltham, MA). Images were deconvolved using a calculated point spread function with the constrained iterative algorithm of Volocity (Perkinelmer Inc, Waltham, MA). Image analysis was carried out using the Quantitation module of Volocity (Perkinelmer Inc, Waltham, MA).

### SIM imaging

Images were acquired using Structured Illumination Microscopy (SIM) performed on an Eclipse Ti inverted microscope equipped with a Nikon Plan Apo TIRF objective (NA 1.49, oil immersion) and an Andor DU-897X-5254 camera. Laser lines 405, 488 and 561 nm were used. Z-step size for Z stacks was set to 0.120 um, which is well within the Nyquist criterion. For each focal plane, 15 images (5 phases, 3 angles) were captured with the NIS-Elements software. SIM image processing and reconstruction were carried out using the N-SIM module of the NIS-Element Advanced Research software. Image analysis was carried out using the Quantitation module of Volocity (Perkinelmer Inc, Waltham, MA) with *x* and *y* binning resolution of 32 nm.

### 3C library preparation

Limbs from ~70 E11.5 embryos, 3 E11.5 embryos with the limbs and heads removed, and the heads of 3 E11.5 embryos were collected in 15 ml tubes with enough PBS to cover them and dissociated. Cells were fixed with 1% formaldehyde for 10 min at room temperature (r.t.). Crosslinking was stopped with 125 mM glycine, for 5 min at r.t. followed by 15 min on ice. Cells were centrifuged at 400 *g* for 10 min at 4°C, supernatants removed and cell pellets flash frozen on dry ice.

Cell pellets were treated as previously described (Dostie and Dekker 2007; Ferraiuolo et al., 2010; Williamson et al., 2014). HindIII-HF (NEB) was the restriction enzyme used to digest the crosslinked DNA.

### 5C primer and library design

5C primers covering the *Usp22* (mm9, chr11: 60,917,307-61,003,268) and *Shh* regions (mm9, chr5: 28,317,087-30,005,000) were designed using ‘my5C.primer’ (Lajoie et al. 2009) and the following parameters: optimal primer length of 30 nt, optimal TM of 65°C, default primer quality parameters (mer:800, U-blast:3, S-blasr:50). Primers were not designed for large (>20 kb) and small (<100 bp) restriction fragments, for low complexity and repetitive sequences, or where there were sequence matches to >1 genomic target. The *Usp22* region was used to assess the success of each 5C experiment but was not used for further data normalization or quantification.

The universal A-key (CCATCTCATCCCTGCGTGTCTCCGACTCAG-(5C-specific)) and the P1-key tails ((5C-specific)-ATCACCGACTGCCCATAGAGAGG) were added to the Forward and Reverse 5C primers, respectively. Reverse 5C primers were phosphorylated at their 5′ ends. An alternating design consisting of 365 primers in the *Shh* region (182 Forward and 183 Reverse primers) was used. Primer sequences are listed in Table S6.

### 5C library preparation

5C libraries were prepared and amplified with the A-key and P1-key primers as described in (Fraser et al. 2012). Briefly, 3C libraries were first titrated by PCR for quality control (single band, absence of primer dimers, etc.), and to verify that contacts were amplified at frequencies similar to that usually obtained from comparable libraries (same DNA amount from the same species and karyotype) (Dostie and Dekker 2007, Dostie, et al. 2007, Fraser, et al. 2010). We used 1 - 10 μg of 3C library per 5C ligation reaction.

5C primer stocks (20 μM) were diluted individually in water on ice, and mixed to a final concentration of 2 nM. Mixed diluted primers (1.7 μl) were combined with 1 μl of annealing buffer (10X NEBuffer 4, New England Biolabs Inc.) on ice in reaction tubes. 1.5 μg salmon testis DNA was added to each tube, followed by the 3C libraries and water to a final volume of 10 μl. Samples were denatured at 95°C for 5 min, and annealed at 55°C for 16 hours. Ligation with Taq DNA ligase (10 U) was performed at 55°C for one hour. One tenth (3 μl) of each ligation was then PCR-amplified individually with primers against the A-key and P1-key primer tails. We used 26 cycles based on dilution series showing linear PCR amplification within that cycle range. The products from 3 to 5 PCR reactions were pooled before purifying the DNA on MinElute columns (Qiagen).

5C libraries were quantified by bioanalyser (Agilent) and diluted to 26 pmol (for Ion PGM™ Sequencing 200 Kit v2.0). One microlitre of diluted 5C library was used for sequencing with an Ion PGM™ Sequencer. Samples were sequenced onto Ion 316™ Chips following the Ion PGM™ Sequencing 200 Kit v2.0 protocols as recommended by the manufacturer (Life Technologies™).

### 5C data analysis

Analysis of the 5C sequencing data was performed as described in (Berlivet et al., 2013). The sequencing data was processed through a Torrent 5C data transformation pipeline on Galaxy (https://main.g2.bx.psu.edu/). Data was normalized by dividing the number of reads of each 5C contact by the total number of reads from the corresponding sequence run. All scales shown correspond to this ratio multiplied by 10^3^. The number of total reads and of used reads is provided for each experiment in Table S7. The unprocessed heatmaps of the normalized 5C datasets can be found in Figure S4. 5C datasets are uploaded to the Gene Expression Omnibus (GEO) website http://www.ncbi.nlm.nih.gov/geo/ accession number: GSE79947.

## Results

### Increased co-localisation of ZRS with *Shh* in the limb ZPA at E10.5 and E11.5

Previous analyses of the chromatin dynamics involved in the long-range regulation of *Shh* by ZRS has produced contradictory results, which could be due to the different temporal stages of development assayed (Amano et al., 2009; Lettice et al., 2014). To resolve this issue we carried out FISH on whole embryo sections that include posterior and anterior forelimb tissue from E10.5, E11.5 and E14.5 developmental stages (Figure 1B). In murine embryos, *Shh* is expressed within the ZPA at the two earlier stages but is switched off in the limb by E14.5 (Riddle et al., 1993). We compared inter-probe distances (Figure S1A shows representative images) and co-localisation frequencies (Figure 1C; left) between the gene and enhancer in tissue across the anterior-posterior axis of the distal forelimb bud. In addition proximal limb tissue and the adjacent flank where *Shh* is not expressed were compared.

By conventional wide-field deconvolution microscopy, the proportion of co-localised (<200 nm) *Shh* and ZRS probe pairs in ZPA cells at stages when *Shh* is expressed (E10.5, E11.5), was significantly higher (35%) than in inactive limb regions and the flank (distal anterior *p* < 0.05, proximal *p* < 0.01, flank *p* < 0.001 (Figure 1C, Table S2)). By E14.5 when *Shh* expression in the limb has ceased, the *Shh*-ZRS co-localisation frequency in distal posterior cells is significantly reduced, compared to E10.5 and E11.5 ZPA (E10.5 *p* = 0.006, E11.5 *p* = 0.01) (Figure 1D). At this later stage, differences in co-localisation frequencies between the distal posterior forelimb region (~20%) and the other limb regions and the flank mesoderm are also no longer detected (Figure 1C).

*Dpp6* is located the same linear genomic distance away from *Shh* as the ZRS, but in the other direction and outside of the *Shh* regulatory domain (Figure 1A). In contrast to the spatial proximity of *Shh*-ZRS, *Shh* and *Dpp6* are predominantly located greater than 400 nm apart (co-localisation frequency < 5%) (Figure 1C; right) for all tissues and developmental stages examined.

The greater co-localisation of the active enhancer (ZRS) with its target gene (*Shh*) in the ZPA at E10.5 is similar to what we reported for these loci at E11.5 (Lettice et al. 2014) and to the co-localisation frequency of *Hoxd13* and its GCR enhancer in distal posterior expressing limb tissue and cell lines at E10.5 (Williamson et al., 2012; Williamson et al., 2014). These differences in chromatin conformation between active and inactive tissues contradicts the previous report suggesting an equivalent rate of *Shh*-ZRS co-localisation on both sides of the distal limb field at this developmental stage (Amano et al., 2009).

### Super-resolution imaging identifies *Shh*/ZRS co-localisation of most alleles in the ZPA

These data are consistent with gene-enhancer co-localisation during long-range regulation. From the images taken by conventional light-microscopy of the tissue sections from the three developmental stages it was apparent that *Shh* and ZRS are consistently very close in the nucleus, with differences in spatial distance frequently down to the signal centroids being in different layers of the *z* stack – the dimension with the lowest spatial resolution in the microscope. We therefore re-analysed the tissue sections containing E10.5 and E11.5 distal anterior and posterior (ZPA) cells by structured illumination microscopy (3D-SIM) (Figure 2A). This technique doubles the resolution limit in all dimensions (Toomre & Bewersdorf, 2010) and has previously been combined with 3D-FISH (Nora et al., 2012; Patel et al., 2013).

**Figure 2.**
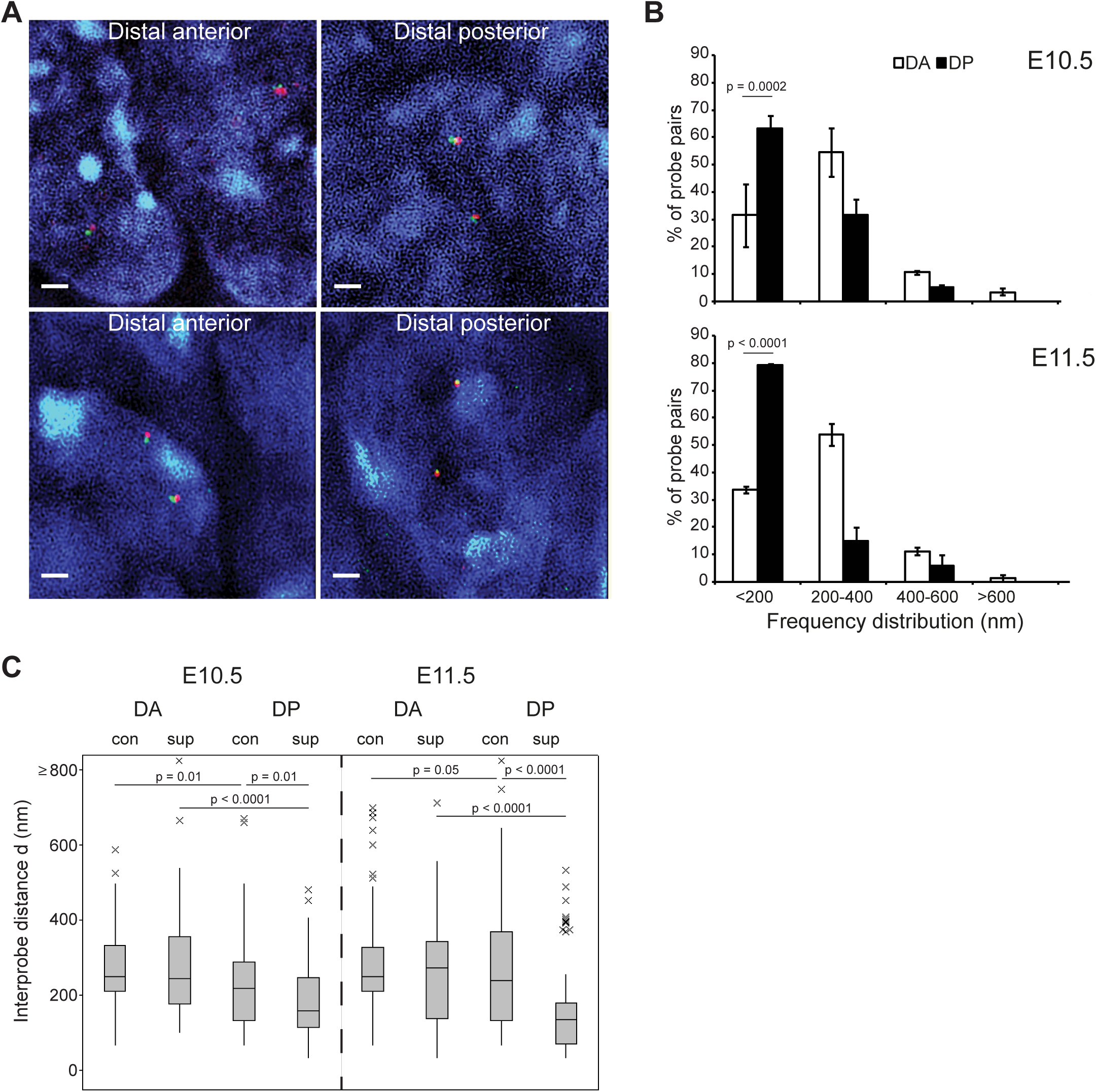
Super-resolution imaging identifies the majority of*Shh*-ZRS probes as co-localised in ZPA tissue. (**A**) Nuclei captured by super-resolution SIM imaging from the distal forelimb of E10.5 and E11.5 embryos after FISH with *Shh* and ZRS probe pairs. Scale bars = 1 μm. (**B**) Frequency distributions of *Shh*-ZRS inter-probe distances (d) measured from SIM images in 200 nm bins, in distal anterior (DA) and distal posterior (DP) regions of the murine forelimb at E10.5 and E11.5. n = 67-100 (alleles). Error bars represent SEM obtained from two different tissue sections from 1 embryo. The statistical significance between data sets was examined by Fisher’s Exact Tests. (**C**) Boxplots show the distribution of *Shh*-ZRS inter-probe distances (d in nm) in E10.5 and E11.5 DA and DP captured by conventional (Con) and structured illumination (SIM) microscopy. Line = median, box = interquartile range, whiskers = 95% range. The statistical significance between data sets was examined by Mann-Whitney U Tests.

The greater resolution afforded by 3D-SIM, particularly for the *z* (depth) dimension (120 nm compared to 200 nm in conventional widefield microscopy), not only confirmed the difference in *Shh*/ZRS co-localisation frequency between ZPA and distal anterior limb bud but also suggests that conventional microscopy does not fully capture the proportion of co-localised *Shh*/ZRS probe pairs, especially in the *Shh*-expressing tissues where it now peaks at 79% (Figure 2B, Table S3). These data suggest that a substantial proportion of *Shh*/ZRS probe pairs with signal centroids not in the same plane of the *z* stack, that have been categorised as adjacent (between 200 nm and 400 nm apart (Figure S1)) due to the low *z* dimension resolution afforded by conventional widefield microscopy, are indeed co-localised in ZPA cells. At both temporal stages the anterior/posterior differences in co-localisation were highly significant (E10.5 *p* = 0.0002, E11.5 *p* = 0.0001). By comparing conventional and SIM data for the *Shh*/ZRS probe pair in anterior and posterior tissues at two developmental stages, we show that median inter-probe distances in distal anterior limb tissues are very similar when measured by either technique (conventional = 250 nm, SIM: E10.5 = 246 nm, E11.5 = 275 nm); whereas, these are significantly different for ZPA cells (E10.5: conventional = 221 nm, SIM = 160 nm, *p* = 0.01; E11.5: conventional = 241 nm, SIM = 136 nm, *p* < 0.0001) (Figure 2C, Tables S4 & S5).

### The *Shh*/ZRS regulatory domain is compact in expressing and non-expressing tissue

Long-range gene/enhancer co-localisation is often depicted as a looping out of the intervening chromatin fibre (Williamson et al., 2011). Our previous work on the HoxD locus implicated a gross compaction of the regulatory region, rather than a simple loop with extrusion of the intervening chromatin, upon activation of *Hoxd13* by the long-range (~250-kb) limb-specific GCR enhancer (Williamson et al., 2012; Williamson et al., 2014). We therefore used 3D FISH and conventional wide-field deconvolution microscopy to measure the spatial distances between either *Shh* or the ZRS, and the SBE4 enhancer that drives *Shh* expression in the forebrain (Figure 3A) (Jeong et al., 2006). SBE4 is located midway through the gene desert separating Shh and ZRS (Figures 1A). If the entire genomic region between the gene and the limb enhancer forms a loop then *Shh*-SBE4 and SBE4-ZRS distances should be greater than those between *Shh* and ZRS.

**Figure 3.**
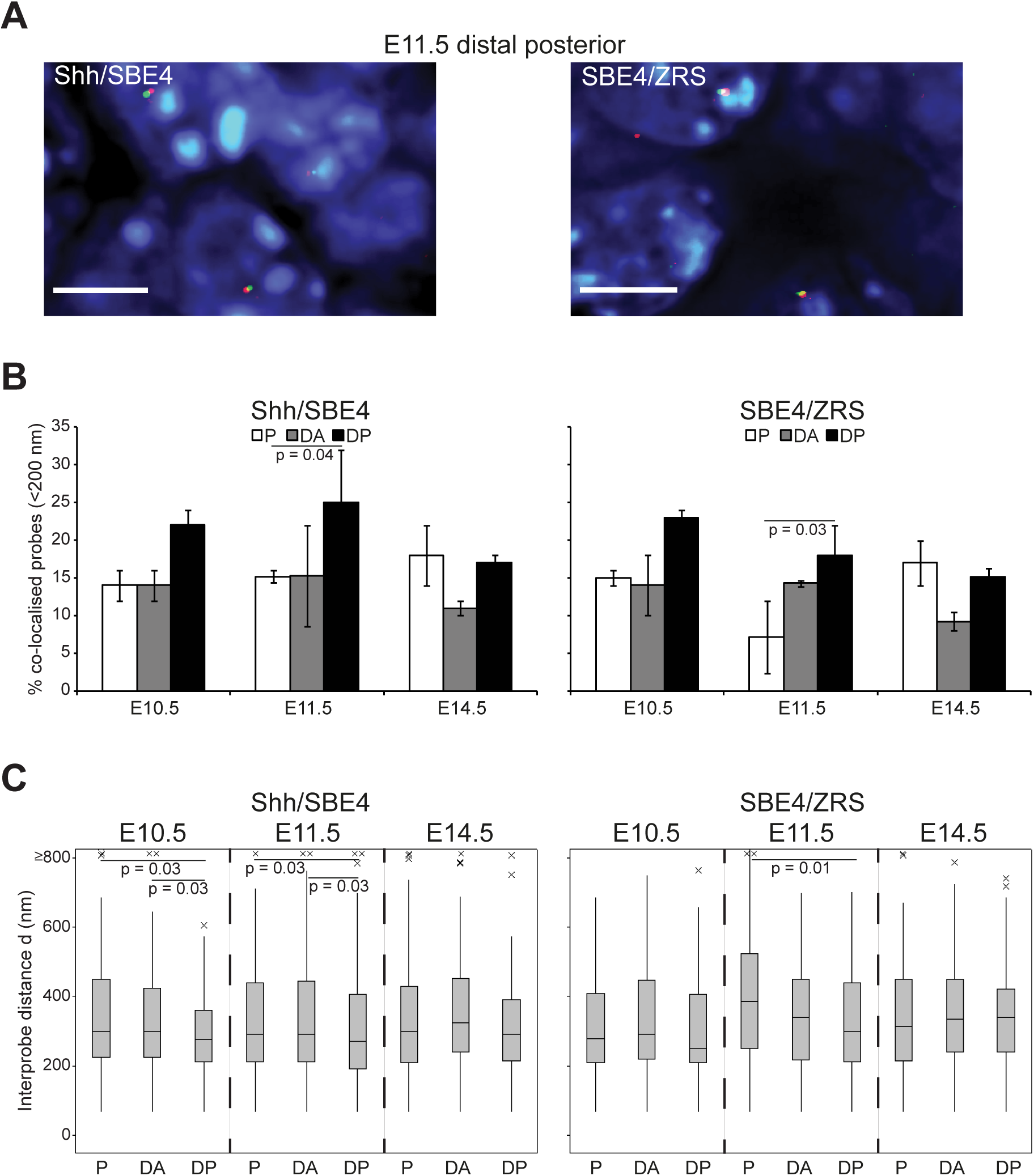
The*Shh*-ZRS regulatory domain is maintained in a compact chromatin conformation in expressing and non-expressing tissue. (**A**) Images of representative nuclei from E11.5 ZPA tissue showing FISH signals for *Shh*/SBE4, SBE4/ZRS probe pairs. Scale bars = 5 μm. (**B**) Comparison of the proportion of co-localised *Shh/SBE4* and SBE4/ZRS probe pairs (<200nm) across the three temporal developmental stages for distal anterior and posterior and proximal forelimb tissue. Error bars represent SEM obtained from two or three different tissue sections from 1-2 embryos. The statistical significance between data sets was examined by Fisher’s Exact Tests. P proximal limb, DA: distal anterior limb, DP: distal posterior limb (ZPA in E10.5 and E11.5 sections). n = 70 – 100 (alleles). (**C**) Boxplots showing the distribution of interprobe distances (d) in nanometres between *Shh*/SBE4 and SBE4/ZRS in E10.5, E1 1.5 and E14.5 distal anterior and posterior and proximal forelimb. The statistical significance between data sets was examined by Mann-Whitney U Tests.

At both temporal stages (E10.5 and E11.5) when *Shh* is active in the distal posterior limb mesenchyme, but not at E14.5, *Shh* is closer to SBE4, and Shh/SBE4 co-localisation frequencies are higher, compared to the other tissues analyzed (Figure 3B & C, S2A & B, and Table S5). These data suggest that the genomic region between *Shh* and the ZRS is folded into a compact chromatin domain, which is at its most compact in distal posterior *Shh*-expressing cells. However, what is also apparent is that the spatial distance between *Shh* and the ZRS is less than both *Shh*-SBE4 and SBE4-ZRS in most expressing and non-expressing tissues (Figure S2C). These differences are significant for most tissues analysed and intriguingly is particularly apparent at E14.5, well past the stage of limb-specific *Shh* activity and therefore could be indicative of a constitutive chromatin conformation.

### Topography of the *Shh* regulatory domain is maintained throughout the E11.5 embryo

Using FISH we could only infer the possibility of a Shh regulatory domain from the spatial relationships of three genomic loci across the *Shh*-ZRS region. To gain a more complete view of genome architecture over the locus, we used 5C to determine the frequency of cross-linked interactions captured between sequences in the ~1.7 Mb region from *Irsig1* ~400kb 3′ of *Shh* to *Ube3c* ~350kb beyond the ZRS (Figure 1A) in dissected limb buds from ~70 E11.5 embryos (x2 replicates) (Figures 4A, left-hand heatmap, S3A and S4). Three interaction domains can be identified; with the middle topologically associated domain (TAD) containing *Shh* and its entire known regulatory elements with the boundaries located 3′ of *Rbm33* and within the 5′ end of *Lmbr1*. This *Shh* regulatory TAD corresponds well with that identified by Hi-C in mouse ESCs (Dixon et al., 2012)

**Figure 4.**
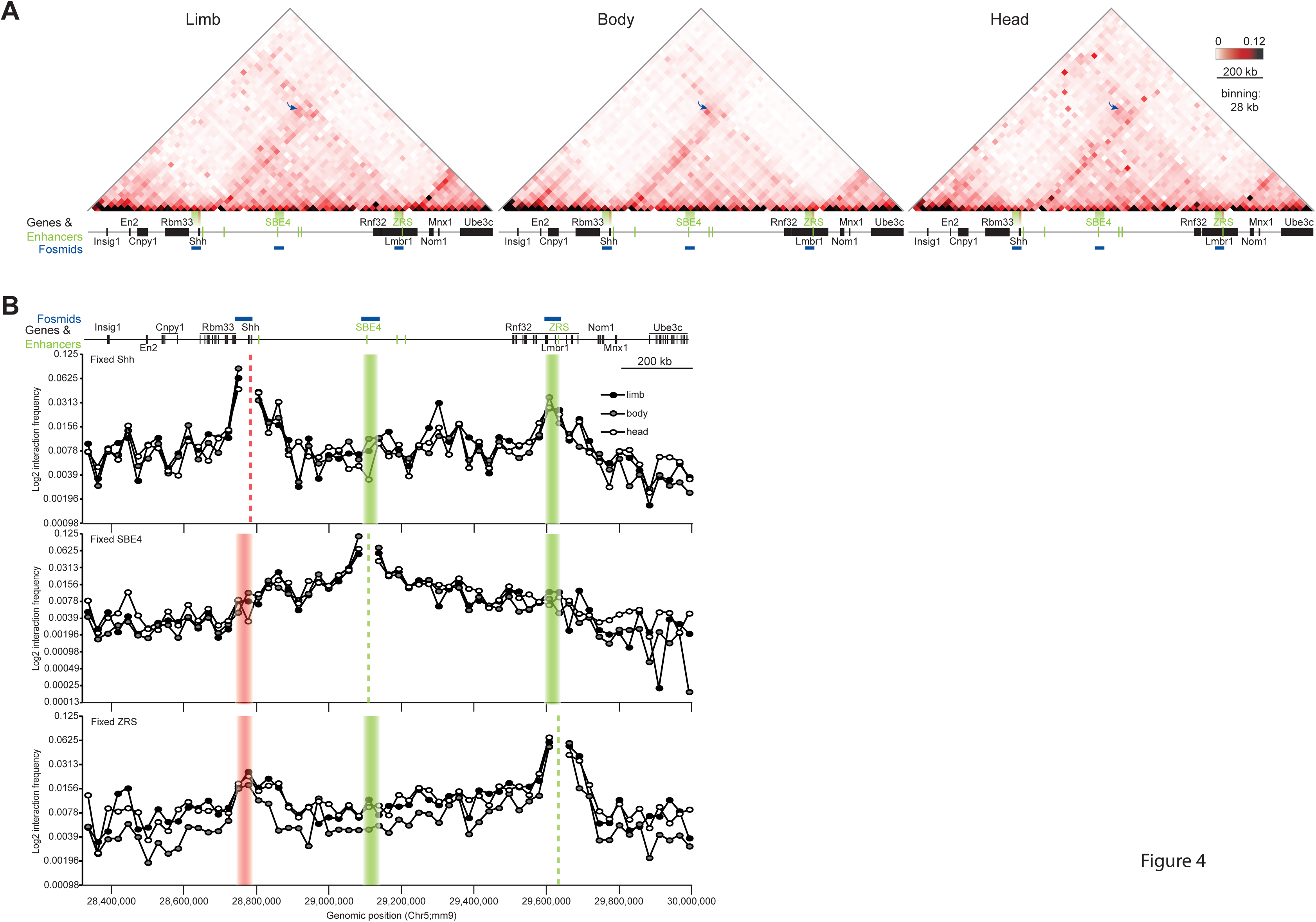
5C-seq identifies enriched interactions between*Shh* and ZRS in E11.5 embryos. (**A**) Heat-maps showing 5C data from cells of the limbs, bodies and heads of E11.5 embryos, across the 1.7-Mb Shh region shown in Figure 1. Heat map intensities represent the average of interaction frequency for each window, colour-coded according to the scale shown. Interaction frequencies were normalized based on the total number of sequence reads in the 5C data set and the data shown is binned over 28-kb windows. Arrows indicate interaction frequencies between windows containing *Shh* and ZRS. Data for biological replicates are in Supplemental Figure S3 A and unprocessed normalized data are shown in Supplemental Figure S4. (**B**) Virtual 4C analysis obtained by extracting 5C interactions with viewpoints fixed at *Shh*, SBE4 and ZRS. Dashed lines indicate the position of the fixed viewpoint from the *Shh* genomic region (orange) or regulatory elements (green). Data from limbs are in closed black circles, bodies is closed grey circles and heads in open circles.

In limb cells 5C cross-linked interactions are enriched between genomic fragments across the *Shh* and ZRS loci (Figures 4A, left-hand heatmap, S3A and S4). The general spatial proximity of *Shh* and the ZRS detected by FISH and inferred from enriched 5C interaction frequencies in expressing and non-expressing tissues suggests that this conformation is constitutive. To determine whether the high cross-linking efficiency of *Shh* and ZRS identified in E11.5 limb buds can also be detected in tissues where the ZRS is not active we carried out 5C on cells derived from the bodies and heads of E11.5 embryos. Even with the vast majority of these cells not expressing *Shh*, high read frequencies between *Shh* and ZRS were captured (Figures 4A, middle and right-hand heatmaps, and S4). Moreover, the same TAD structures can be discerned across the genomic region analysed.

To examine more closely the regions probed by FISH (*Shh*, SBE4 and ZRS) we generated “virtual 4C” plots from the 5C data (Figures 4B, S3B) (Williamson et al., 2014). From the viewpoint of *Shh*, overall interaction frequencies with the rest of its regulatory domain is similar in limb-, body-and head-derived tissues, and are not substantially higher than those extending into the adjacent TAD 3′ of *Shh* (Figures 4B compare the top track with the track that profiles SBE4 located in the middle of a TAD). Highest interaction frequencies for *Shh*, apart from genomic regions immediately adjacent, are with regions within the neighbourhood of ZRS (limb-specific high interactions with a loci within the gene desert that does not contain any known regulatory elements was not identified in the limb replicate data (Figure S3B)). ZRS has reciprocal enriched interactions with the *Shh* region (Figures 4B, bottom track). Therefore, virtual 4C plots from all three locations of E11.5 embryos analysed have largely corresponding interaction frequencies of the pseudobaits with the rest of the genomic region, highlighting the specificity of cross-linking frequency between *Shh* and ZRS.

## Discussion

### Activation of *Shh* in the limb bud is accompanied by co-localisation with the ZRS

Using 3D-FISH and super-resolution imaging, we provide compelling evidence that co-localisation (<200 nm) between *Shh* and the ZRS enhancer is associated with *Shh* expression in the ZPA region of the distal posterior forelimb bud, to an extent not seen in control tissues, including the limb bud after Shh expression has ceased at E14.5 (Figures 1 and 2). The co-localisation frequencies detected by super-resolution microscopy rise to almost 80% at E11.5 – suggesting that the vast majority of Shh alleles in the ZPA are juxtaposed to the ZRS located 1Mb of genomic distance away. Analysis of the FISH images by either conventional wide-field, or structure illumination microscopy, showed a significantly higher gene-enhancer co-localisation frequency in the ZPA than in nuclei from the distal anterior region of the same limb buds (Figures 1 and 2). This anterior-posterior difference in chromain folding is consistent with our previous analysis for *Shh*-ZRS in E11.5 fore-and hindlimbs (Lettice et al. 2014) and is similar to the preferential co-localisation of *Hoxd13*-GCR in E10.5 distal posterior limb buds (Williamson et al., 2012). Like *Shh*, *Hoxd13* expression is restricted to the posterior margin of the distal limb bud at this stage. These data contradict previously published work that could not identify any signficant difference in *Shh*-ZRS proximity between the *Shh*-expressing ZPA and distal anterior cells (Amano et al., 2009). These data were derived from single cell suspensions of dissected tissue from specific points across the distal limb bud whereas our data are from sections cut through whole embryos; therefore cell/tissue preparation may be a factor in discrepencies between the data sets. On the other hand, both data sets identify a clear difference in co-localisation between ZPA cells and cells that never express *Shh*.

### *Shh* and its regulatory elements are located within a compact chromatin domain

Long-range interactions between genes and *cis*-regulatory elements are usually described as loops, often visualized as a coming together of the two loci to the exclusion of the intervening chromatin (Williamson et al. 2011; Fraser et al. 2015). Indeed a looping mechanism in this distal limb could be inferred from the shorter inter-probe distances between *Shh* and ZRS, than for either of these probes with the forebrain SBE4 enhancer – even though the latter is located midway between *Shh* and ZRS on the linear chromosome (Figure S2C). However, the *Shh*-ZRS distances are shorter than distances to SBE4 not only in the ZPA tissue but also in anterior limb and in E14.5 tissues when the ZRS is no longer active. But, *Shh*-ZRS co-localisation frequencies are not significant in these tissues. Another interpretation of the data is that the *Shh* regulatory domain (Figures 4A, S3A, S4) is maintained in a tightly folded chromatin conformation where *Shh* and the ZRS are generally proximal in nuclear space. That this *Shh*-containing TAD is indeed compact can be discerned from the frequency distribution graphs which show that most *Shh*/ZRS, *Shh*/SBE4 and SBE4/ZRS probe pairs are adjacent (200 – 400 nm) or co-localised (<200 nm), with median interprobe distances of between 220 – 345 nm for most tissues and developmental stages (Figures 1B, S2A & B; Table S5).

Moreover, 5C analysis of E11.5 limb bud, body and head cells suggests that the *Shh* regulatory region forms a constitutive self-interacting domain with somewhat enriched interactions between cross-linked DNA fragments from the genomic regions containing *Shh* and ZRS (Figures 4, S3, and S4). This *Shh* TAD has also been identified in ES cells by Hi-C (Dixon et al. 2012).

### Facilitating gene regulation by enhancer – promoter proximity

If the physical interaction of active enhancers and their target gene promoters is essentially a stochastic process, their constitutive relative proximity within the same chromatin domain could be advantageous – for example by reducing the search space of the enhancer for the promoter (Williamson et al., 2011; Benabdallah and Bickmore, 2015). Consistent with this model, we have previously shown that expression levels in the limb of a reporter gene, inserted into several positions across the whole *Shh* regulatory domain, is highest when the reporter inserts close to either the ZRS or *Shh* compared to insertion sites within the intervening gene desert (Anderson et al. 2014).

## Acknowledgements

We thank the staff of the IGMM imaging facility and technical services for their assistance with imaging and sequencing. This work was supported by the Medical Research Council, UK.

